# Development of SSR markers related to seed storability traits in maize subjected to artificial seed aging conditions

**DOI:** 10.1101/833111

**Authors:** Xiaoyang Guo, Chengxu Gong, Shan Liu, Chenchen Feng, Xiao Han, Tingting Lv, Xiaohui Sun, Xiuwei Yang, Yu Zhou, Zhenhua Wang, Hong Di

**Author notes:** contributed equally to this research (Xiaoyang Guo, Chengxu Gong). Corresponding author: Hong Di.

## Abstract

Seed storability is an important and complex agronomic trait in maize because annual seed production considerably exceeds consumption. The viability of seeds decreases over time, even when stored at low temperature, until seeds finally lose viability. In our previous study, two inbred lines with significantly different storability, Dong156 with high storage tolerance and Dong237 with low storage tolerance, were selected over six years using a natural seed aging test. In the present study, an F2:3 population and a RIL (recombinant inbred line) population were constructed from these two inbred lines and used to map QTL (quantitative trait loci) with SSR (simple sequence repeat) markers. A phenotypic index of traits related to seed storability that includes germination rate, germination potential, a germination index, a vigor index, seedling weight, and seedling length was generated using the results of an artificial aging treatment. Two consistent regions, *cQTL-*7 on chromosome 7 and *cQTL-*10 on chromosome 10, were identified by comparing QTL analysis results from these two populations. After genotyping SSR markers in these two regions, *cQTL-*7 was remapped to between umc1671 and phi328175 in a 7.97-Mb region, and *cQTL-*10 was remapped to between umc1648 and phi050 in a 39.15-Mb region. Four SSR markers linked to *cQTL-*7 and *cQTL-*10, including umc1671, phi328175, umc1648, and phi050, were identified using a Chi-squared test. The combined selection efficiency of these four markers was 83.94% in 85 RIL lines with high storability, and marker umc1648 exhibited the highest efficiency value of 88.89%. These results indicated that the four SSR markers developed in this study could be used for selection of maize germplasm with high seed storability.

## Background

Maize is cultivated worldwide in a range of agroecological environments for food, livestock feed, and as an industrial feedstock [1]. Maize seed must be stored from harvest until the next planting season and sometimes much longer because seed production usually considerably exceeds consumption each year [2]. Seed storability, also known as the longevity of seeds after storage, is an important aspect of seed quality that is inherited polygenically and influenced by the environment [3].

Over time, seeds lose viability due to seed aging or deterioration even when stored at low temperatures. Seed storability is likely to be controlled by multiple genes, and is related to physiological, cellular, biochemical, and metabolic activities. Reactive oxygen species (ROS) that accumulate in seeds during storage oxidize lipids and proteins, and consequently modify the structure and function of membranes [4]. Lipid peroxidation results in high concentrations of H_2_O_2_ and malondialdehyde (MDA), which have been considered critical causes of seed deterioration [5-7]. Other factors have also been implicated in seed storability, including DNA and RNA degradation, mitochondrial activity, and redox homeostasis states [8-10].

To further understand the genetic mechanisms controlling seed storability, researchers have performed proteomic analyses after artificial seed aging treatments in *Arabidopsis*, maize, and rice [11-14].

Proteomic analyses have revealed that loss of seed vigor can be accounted for by protein changes in the dry seeds of *Arabidopsis* [11]. Compared to inviable seeds, the expression of small heat shock proteins, late embryogenesis abundant (LEA) proteins, and antioxidant enzymes increase strongly in high-viability seeds of maize hybrid Zhendan958 [12]. In maize cultivar Dabaitou, embryos from seeds that were artificially aged at 50°C for 5 or 13 d, a total of 40 proteins with changed expression were identified. Artificial ageing increased the expression of proteases that degraded stored proteins, impaired metabolism, and energy supply, and ultimately resulted in seed deterioration [13].

Several redox regulation proteins (mainly glutathione-related proteins) and some disease/defense-related proteins (proteins related to repair or tolerance of DNA-damage, and a putative late embryogenesis abundant protein) also might be correlated with seed storability [14].

Quantitative trait loci (QTL) analysis is a powerful tool for deciphering the molecular basis of complex traits. QTL in several species have been linkage mapped after subjecting seeds to natural or artificial conditions that result in seed aging [15-17]. In soybean, 34 QTL affecting seed storability were identified on 11 chromosomes in two RIL populations. Twenty-one of those QTL clustered into five regions on four chromosomes carrying many QTL and another 13 QTL for seed storability were detected in two other QTL-rich regions [18]. In rice treated with both natural and artificial aging conditions, 13 QTL affecting seed storability were identified on eight rice chromosomes. Two of these QTL were detected more than once under both natural and artificial aging treatments. Four other QTL were detected once under natural aging conditions and seven QTL were detected once under artificial aging conditions [19].

In maize, using an F2:3 population and a RIL population derived from the inbred lines X178 and I178, 13 QTL for the six traits related to seed vigor above were mapped into five chromosome regions [20]. Han et al (2018) detected 74 QTL and 20 meta-QTL (mQTL) related to seed vigor in two related RIL populations derived from the crosses Yu82 × Shen137 and Yu537A × Shen137 subjected to three aging treatments. The chromosome regions containing four key mQTL (*mQTL2-2, mQTL5-3, mQTL6*, and *mQTL8*) located near other important QTL might represent regions containing many QTL for traits associated with seed storability [21]. Earlier, Han et al. (2014) used single-nucleotide polymorphism (SNP) markers to detect 65 QTL for four seed vigor-related traits in two RIL populations derived from the same crosses under various artificial seed aging conditions. They also identified another 18 meta-QTL associated with at least two QTL initially across two populations [22].

Maize seed storability is a seriously under-studied problem considering its importance. Our previous study demonstrated that two maize inbred lines bred by our group, Dong156 and Dong237, showed significant differences in seed storability under natural storage conditions. In the present study, an F2:3 and RIL population derived from these two lines were subjected to an artificial aging treatment to (1) map QTL for traits related to seed storability, (2) examine the consistency of QTL regions across the two populations, and (3) develop and verify SSR markers located in the consistent QTL.

## Materials and methods

### Plant materials

The two inbred lines used in the present study, Dong156 and Dong237, were developed at Northeast Agricultural University (NEAU) in Harbin, China, and have good and poor seed storability, respectively. A total of 300 F2 progeny were produced from the F1 progeny of Dong156 × Dong237. The F2:3 population was grown under typical maize cultivation conditions at the NEAU experimental farm in Harbin, Heilongjiang Province (N45°46′ and E126°54′ situated at an elevation of 126 m above mean sea level) along with the parental lines in 2012. We harvested seeds from 267 F3 ears on F2 plants. F2 plants were genotyped with SSR markers, and F3 seeds from each F2 plant were phenotyped for seed storability. A RIL population consisting of 288 lines was developed from five successive generations of self-pollination of F2 individuals in 2016. The F7 RIL lines were then genotyped using SSR markers, and the F8 seeds harvested were utilized for measuring the phenotype. All of the harvested seeds were dried in the sun and then stored at −20°C before testing seed storability under an artificial aging treatment.

Both parental lines, 267 F2 individuals, and the 288 RIL lines were planted in three-row plots from April to September in 2012 and 2016, respectively. Each plot was 3.0 m in length and 0.65 m wide, with an interval of 0.20 m between plants. Experimental fields were managed in a manner consistent with regional commercial maize production practices.

### SSR assays

Genomic DNA was extracted from the two fresh youngest leaves of 20-day-old plants using the CTAB protocol [23]. We then genotyped 1349 SSR markers distributed on the 10 chromosomes of maize (MaizeGDB (http://www.maizegdb.org/) in the parental inbred lines Dong156 and Dong237. Markers showing clear polymorphism between the parental lines were used to genotype individuals in the F2 and RIL populations in different years. SSR analyses including PCR, polyacrylamide gel electrophoresis, and silver staining were performed following the protocols described by Di et al (2015) [24].

### Artificial seed aging treatment

Seeds were sterilized in a solution of 10% sodium hypochlorite in water for 20 min and then washed three times in deionized water. We performed an artificial aging germination (AG) treatment as follows: 50 seeds were enclosed in a mesh bag, which was then placed in a thermostatic water bath at 58 °C and 100% relative humidity for 0 h or 1 h. The 0 h treatment was used as a control. After the artificial aging treatment, seeds were dried at room temperature (∼25 °C) for 2–3 d. We then performed a germination test in a bed of sand following the standard procedures of the International Seed Testing Association [25], with three independent replications per line. Germination was recorded each day and used to calculate the germination index (GI). GI=∑Gt/Dt, where Dt is the number of days after incubation at 58 °C, and Gt is the number of seeds germinated on that day. After holding seeds at 25°C for 4 d, germination energy (GE) was calculated as the proportion of seeds that had germinated relative to the total number of seeds tested. At 7 d, germination percentage (GP) was calculated as the percentage of normal seedlings to the total number of seeds tested. Roots were then removed from the seedlings, and seedling length (SL) and seedling weight (SW) were measured. A vigor index (VI) was then calculated as VI = GP × SW and a simple vigor index (SVI) was calculated as SVI: VI = GI × SW. The relative aging indices were calculated as the ratio of the above values of GI, GE, GP, SL, VI, and SVI for the artificially aged group to those of the control group. Data analyses were performed using SPSS 16.0 and SPSS 19.0 software (IBM Corp., Armonk, NY, USA) (https://www.ibm.com/analytics/spss-statistics-software) [26,27], and Microsoft Excel 2010 [28]. Multiple comparisons were performed using Student’s *t-*test (GraphPad Prism Software, La Jolla, CA, USA). All values were expressed as mean ± SD. Differences with a *p*-value < 0.05 were considered statistically significant.

### Linkage mapping and QTL analysis

Comparing the phenotypic and genotypic data, QTL were detected by composite interval mapping using QTL analysis software IciMapping 4.0 and 4.1 [29] separately at a LOD threshold above 3.0. We calculated the LOD threshold with 1000 permutations P = 0.05 and a 5-cM window [30]. A *t*-test was used for multiple comparisons.

Additional polymorphic SSR markers were added to the regions containing consistent QTL detected in the two populations, and QTL related to seed storability were reanalyzed.

### SSR marker development and genotyping

Thirty lines with very good or poor seed storability where chosen from the RIL population to create a pair of mixed pools, and the genotypes of the two parental lines and two pools were detected using the SSR markers linked to the consistent QTL. Chi-squared tests were conducted to analyze the genotyping results, and SSR markers that were polymorphic between the two parental lines and two pools were chosen. Markers were identified that could detect genotypes with good seed storability phenotypes in 85 RIL lines with good seed storability.

## Results

### Phenotypic performance for seed storability-related traits in the parental lines, F2:3, and RIL populations

Dong156 and Dong237 exhibited significant differences (P <0.01) in RGE (relative germination energy), RGP (relative germination percentage), RGI (relative germination index), RVI (relative vigor index), and RSVI ((relative simple vigor index) after the artificial aging treatment, and significant differences (P <0.05) in relative seedling length (RSL) (Table1). The values of each of the compared traits in Dong156 exceeded those in Dong237. These results indicated highly dominant differences in seed storability between Dong156 and Dong237 and that these lines would be suitable for creating a segregating population for genetic analysis.

**Table 1.** Statistical analysis of indices related to seed storability in RIL population.

| Seed Treatment | Trait | Dong156 | Dong237 | Mean ±SD | Range | S | CV (%) | Skewness | Kurtosis |
| --- | --- | --- | --- | --- | --- | --- | --- | --- | --- |
| Control | GP | 100.00 aA | 98.00 aA | 92.86±0.35 | 48.00-100.00 | 8.43 | 9.08 | -2.346 | 7.570 |
| | GE | 86.00 aA | 84.00 aA | 86.78±0.51 | 30.00-100.00 | 12.33 | 14.21 | -1.983 | 4.991 |
| | GI | 35.96 aA | 35.40 aA | 34.28±0.15 | 15.01-37.98 | 3.62 | 10.56 | -2.047 | 5.707 |
| | VI | 1324.06 aA | 1214.76 aA | 1054.34±7.59 | 387.91-1671.33 | 182.11 | 17.27 | -0.369 | 1.031 |
| | SL | 36.81 aA | 34.29 aA | 30.64±0.15 | 21.67-44.01 | 3.47 | 11.33 | 0.288 | 0.516 |
| | SVI | 36.81 aA | 33.63 aA | 28.52±0.19 | 12.40-44.01 | 4.49 | 15.74 | -0.322 | 1.140 |
| Artificial Aging | GP | 92.00 aA | 4.00 bB | 48.40±0.73 | 2.00-96.00 | 21.46 | 44.34 | 0.152 | -0.565 |
| | GE | 49.33 aA | 3.33 bB | 39.98±0.66 | 0.00-90.00 | 19.40 | 48.52 | 0.437 | -0.100 |
| | GI | 28.70 aA | 1.31 bB | 16.53±0.26 | 0.31-35.14 | 7.73 | 46.76 | 0.289 | -0.350 |
| | VI | 682.62 aA | 23.61 bB | 362.18±7.42 | 1.62-982.30 | 218.08 | 60.21 | 0.566 | -0.335 |
| | SL | 23.78 aA | 17.97 bB | 20.61±0.18 | 4.23-32.34 | 5.25 | 25.47 | -0.443 | -0.197 |
| | SVI | 21.88 aA | 0.72 bB | 10.58±0.21 | 0.10-26.90 | 6.12 | 57.84 | 0.444 | -0.595 |
| Relative value | RGP | 92.00 aA | 4.07 bB | 51.91±1.29 | 2.15-97.92 | 21.84 | 42.07 | 0.057 | 0.564 |
| | RGE | 57.42 aA | 3.93 bB | 46.17±1.26 | 0.84-97.80 | 21.29 | 46.11 | 0.315 | 0.194 |
| | RGI | 79.84 aA | 3.69 bB | 48.05±1.26 | 1.58-97.50 | 20.31 | 42.27 | 0.186 | 0.371 |
| | RVI | 51.76 aA | 1.91 bB | 34.10±1.15 | 0.36-92.28 | 19.46 | 57.07 | 0.475 | 0.361 |
| | RSL | 64.77 aA | 52.51 aB | 67.38±0.95 | 15.97-99.58 | 16.09 | 23.88 | -0.480 | 0.067 |
| | RSVI | 59.63 aA | 2.13 bB | 36.68±1.19 | 0.49-93.76 | 20.17 | 54.99 | 0.382 | 0.581 |

The seed germination rate (GR) of the F2:3 population was above 99% before applying the artificial aging treatment. Significant differences were detected among the F2:3 lines for all traits tested. We found that the indices related to seed storability GI, GE, GP, SL, and SVI are normally distributed with transgressive segregation, except for index VI, as expected for patterns of variability in quantitative traits (Figure 1).

**Fig. 1.**
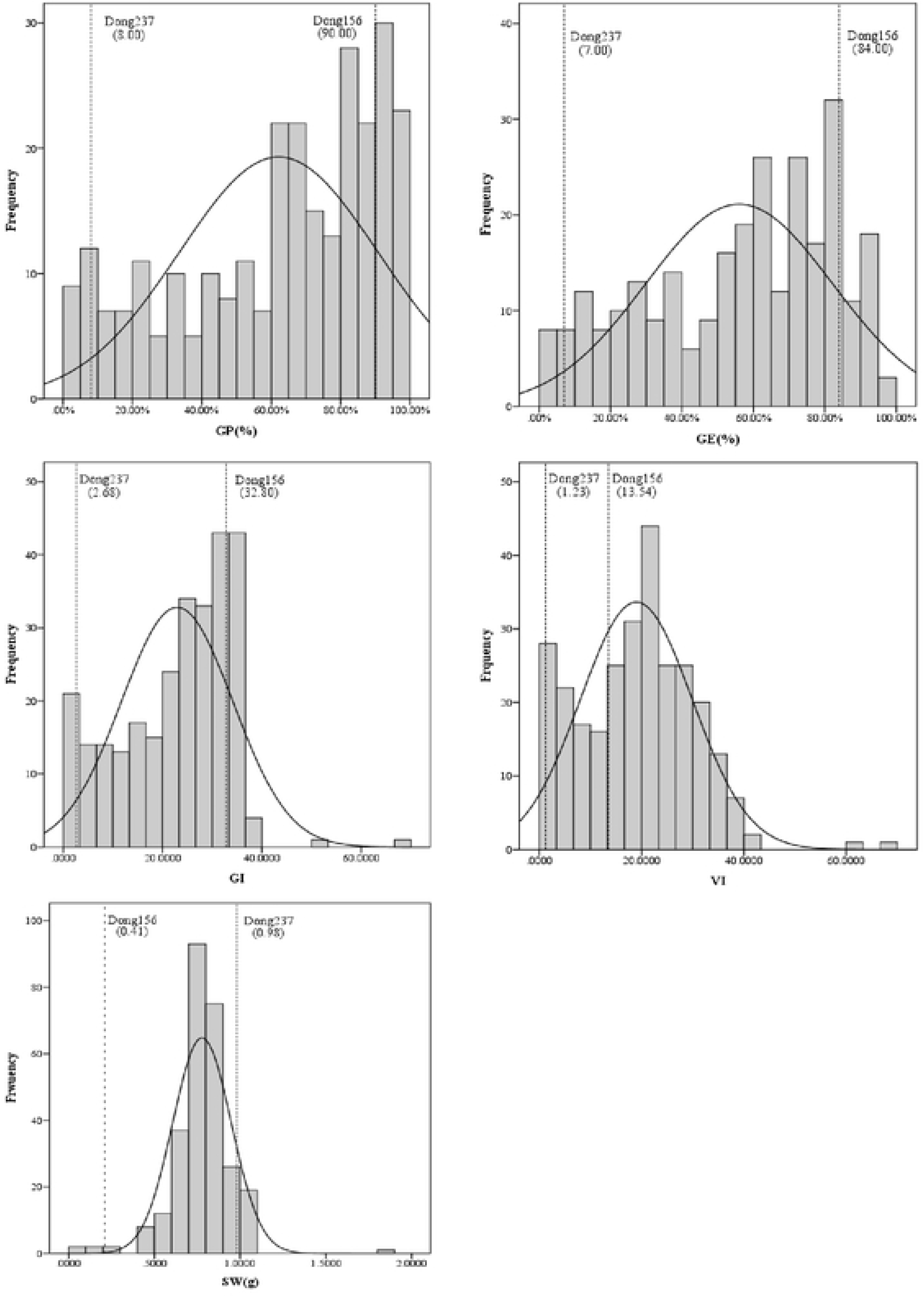
Distribution of seed storability-related traits in an F2:3 population and their parental lines Dong156 and Dong237. GP: germination percentage; GE: germination energy; GI: germination index; VI: vigor index; SW: seedling weight

A great range in variability and significant differences became apparent among the traits RGE, RGP, RGI, RVI, RSVI, and RSL related to seed storability during the standard germination test in the RIL population of 288 lines before and after the artificial aging treatment. Variance coefficients for RGE, RGP, RGI, RVI, and RSVI exceeded 40% except for RSL, with the greatest value (57.07%) for RVI. Analysis of the frequency distributions revealed normal distributions for the relative values of GE, GP, GI, VI, SVI, and SL, as presented in Figure 2 and Table 1. There are also significant differences in GR of RIL lines 7 d after artificial aging (Figure 3).

**Fig. 2.**
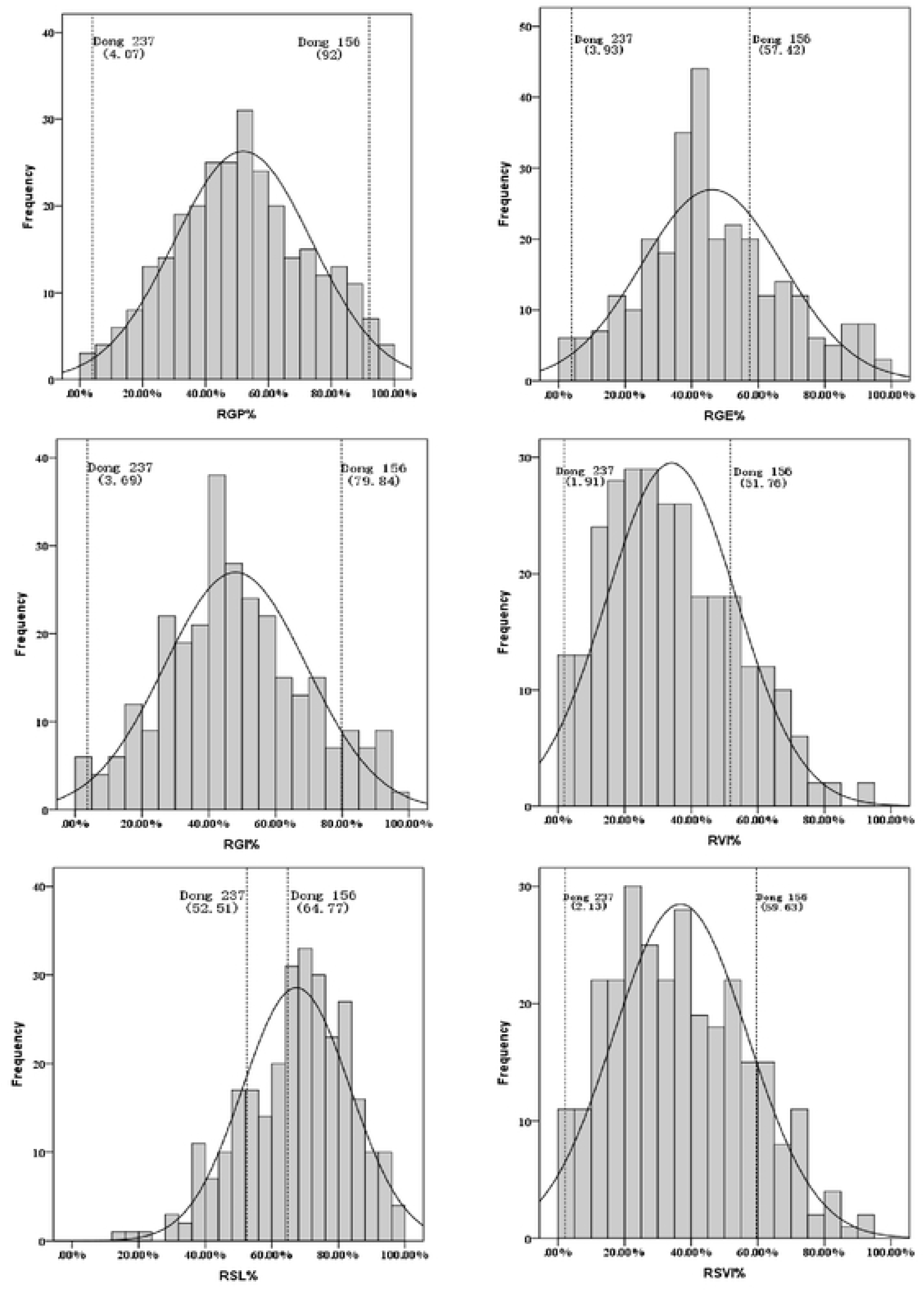
Distribution of relative values for seed storability traits in a RIL population derived from Dong156 × Dong237 and parental lines Dong156 and Dong237. RGP: relative germination percentage; RGE: relative germination energy; RGI: relative germination index; RVI: relative vigor index; RSL: relative seed ling length; RSVI: relative simple vigor index

**Fig. 3.**
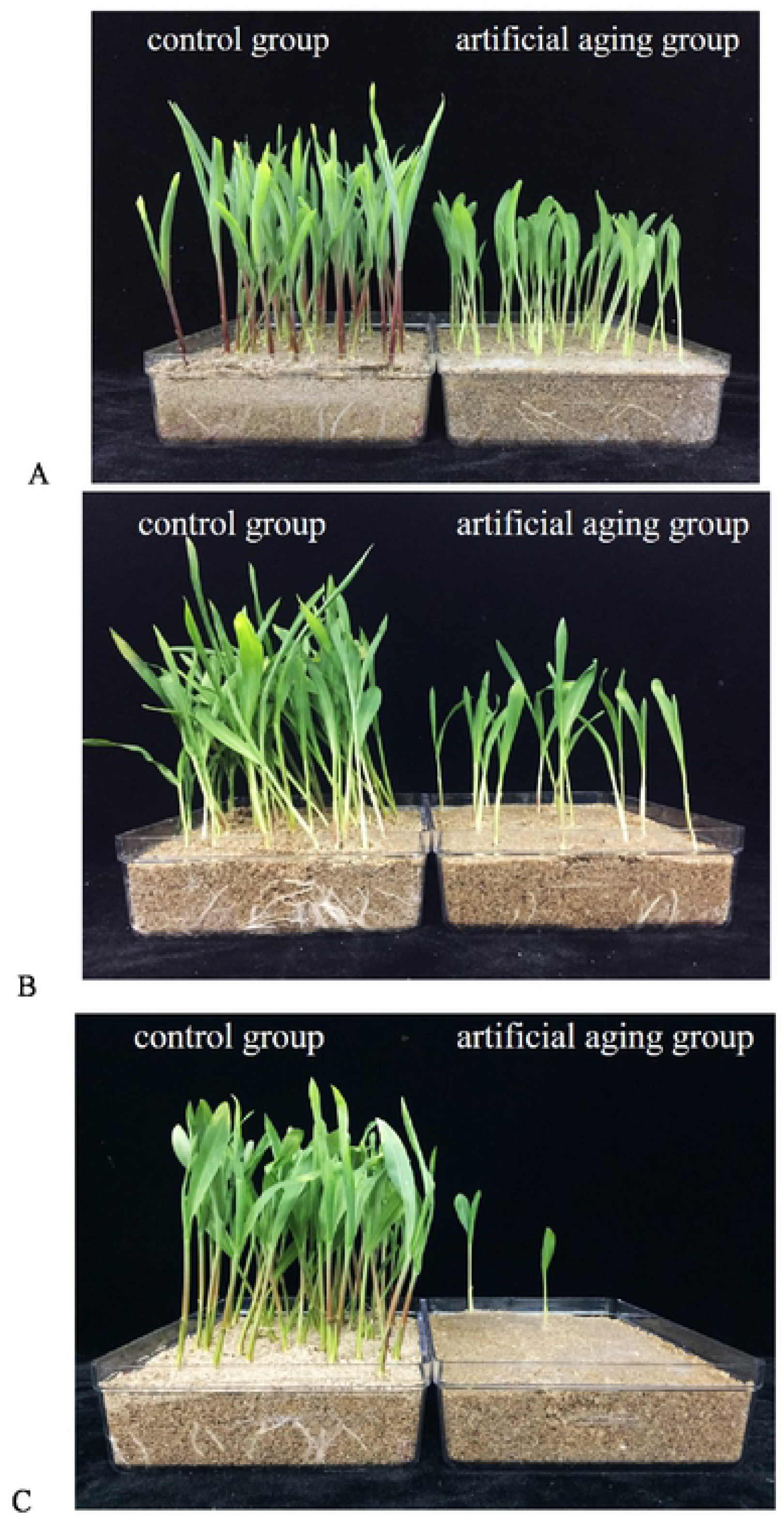
Germination of parental lines Dong156 and Dong237and a RIL line 201-7 derived from Dongl56 × Dong237 at 7 d after treatment in control and artificial seed aging treatment groups. A: parental line Dongl56 with good storability; B: RIL line 201-7 with medium storability; C: parental line Dong237 with poor storability

### Linkage map construction and QTL analysis

The order and positions of 192 polymorphic SSRs in our molecular linkage map for the F2:3 population agreed with those at the MaizeGDB (http://www.maizegdb.org/ssr.php). We detected a total of 11 QTL for seed storability on maize chromosomes 1, 2, 4, 5, 7, 8, and 10 (Fig. 4) in the F2:3 population that was subjected to the artificial aging treatment. The alleles conferred by Dong156 had positive additive effects, thereby increasing seed storability for all QTL detected. *qSVI-10*, which is located between bnlg162 and umc2043 on chromosome 10, explains 24.35% of the phenotypic variation in SVI in this F2:3 population, with a LOD score of 5.34. Between markers umc2063 and umc2400 on chromosome 5, *qSW-5* explains 22.68% of the phenotypic variation in SW in this F2:3 population, with a LOD score of 3.31. We also found that *qSVI-7-2* between markers umc1545 and umc2333 on chromosome 7 explains 20.87% of the phenotypic variation in SVI in this F2:3 population, with a LOD score of 2.63 (Table 2).

**Table 2.**
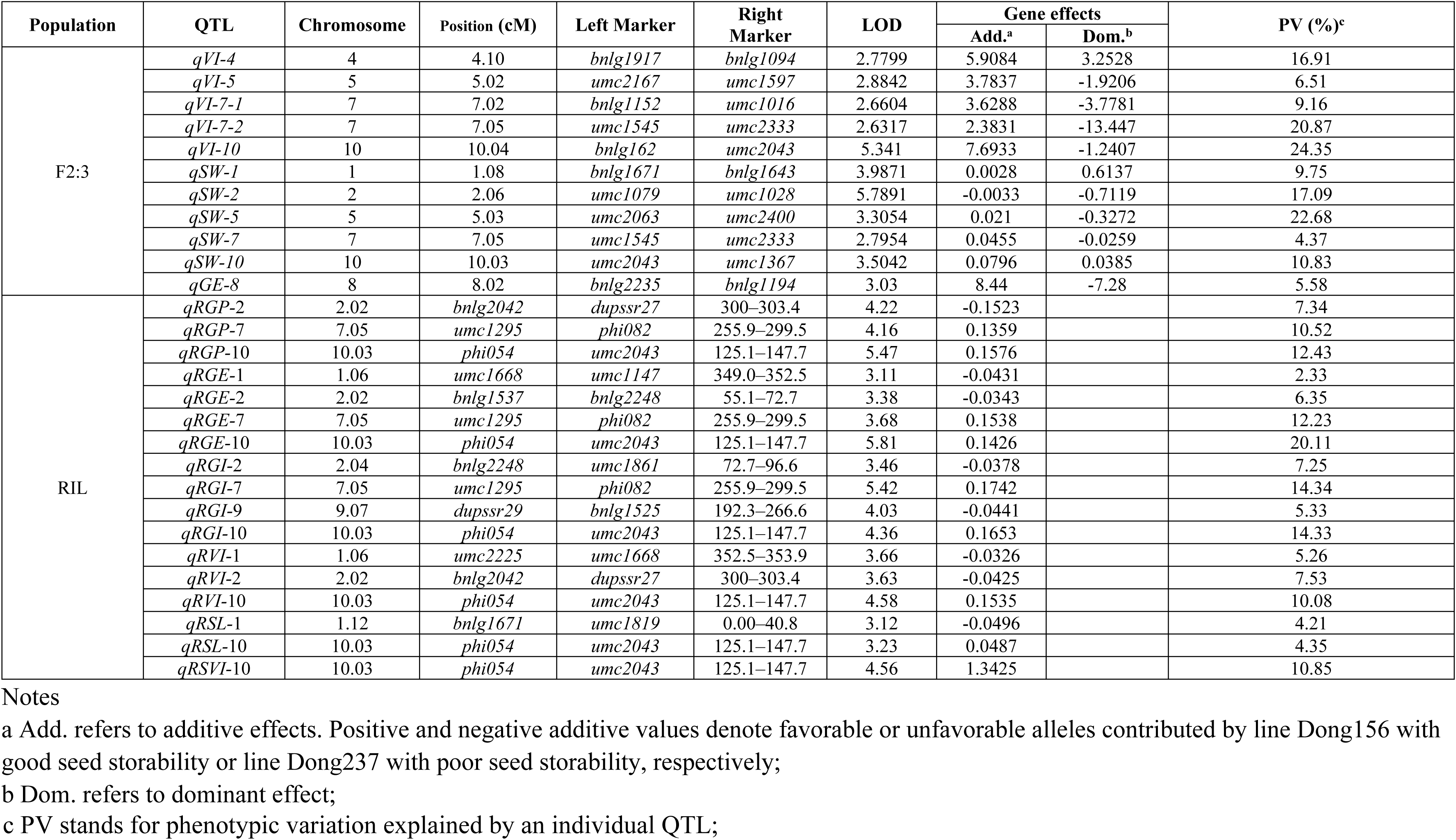
Major QTL for seed storability in F_2:3_ and RIL populations derived from Dong156 and Dong237.

**Fig. 4.**
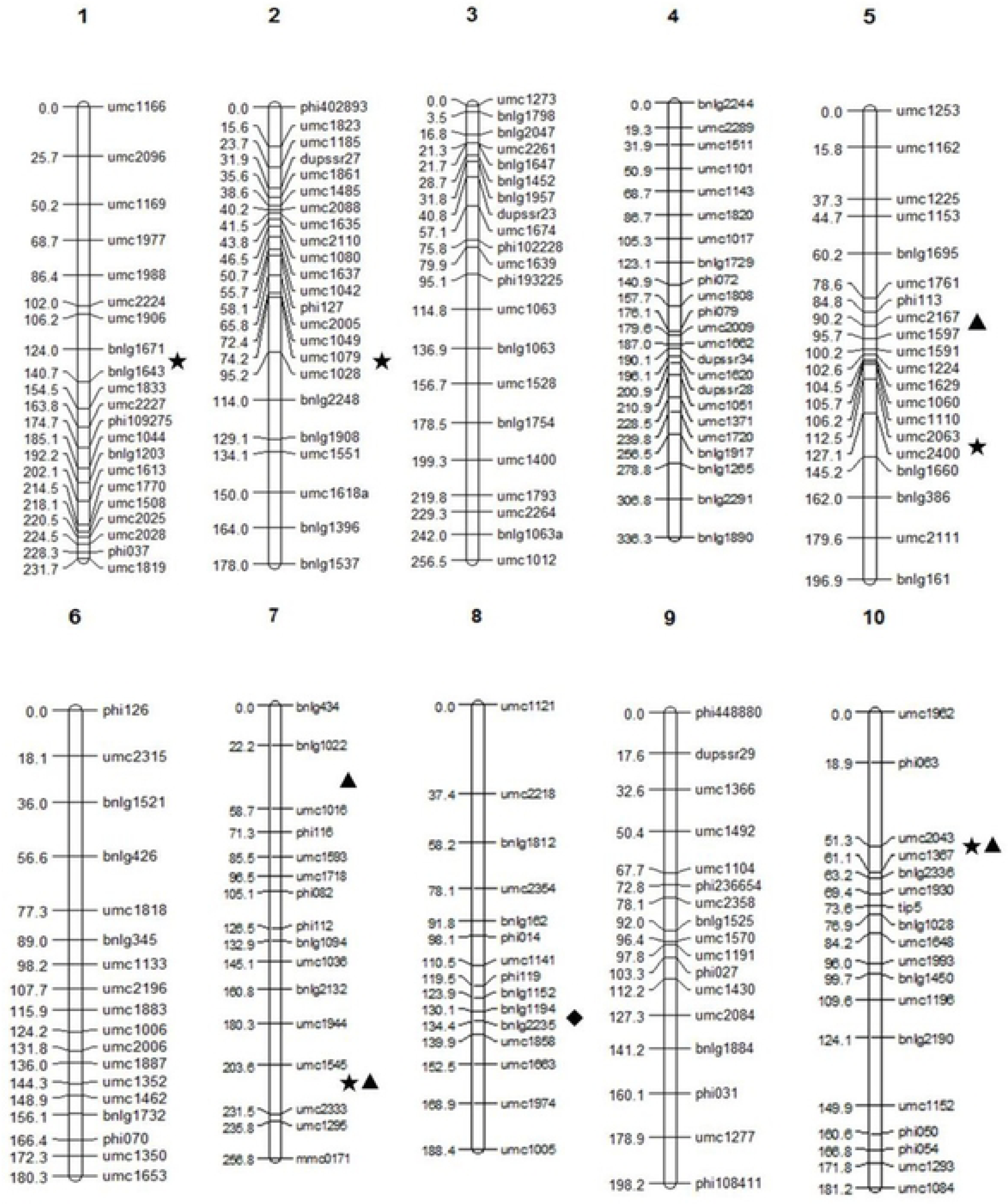
QTL for seed storability related traits on the F2:3 linkage map. **★** refers to locus controlling seedling weight; **▴** refers to locus controlling vigor index; ♦ refers to locus controlling germination energy

The molecular linkage map for the RIL population includes 226 polymorphic SSRs, and with an average spacing of 15.34 cM. Analysis of the phenotypes and genotypes after artificial seed aging in the RIL population revealed 17 QTL affecting seed storability located on chromosomes 1, 2, 7, 9, and 10 (Figure 5) that explain between 2.33 and 20.11% of the phenotypic variation observed in RGE, RGP, RGI, RVI, RSVI, and RSL in the RIL population. Finally, *qRGE-*10, which is located between phi054 and umc2043 on chromosome 10, explains 20.11% of the phenotypic variation in RGE in the RIL population with a LOD score of 5.81 (Table 2). Alleles that increase seed storability for nine QTL detected in the RIL population were conferred from Dong156.

**Fig. 5.**
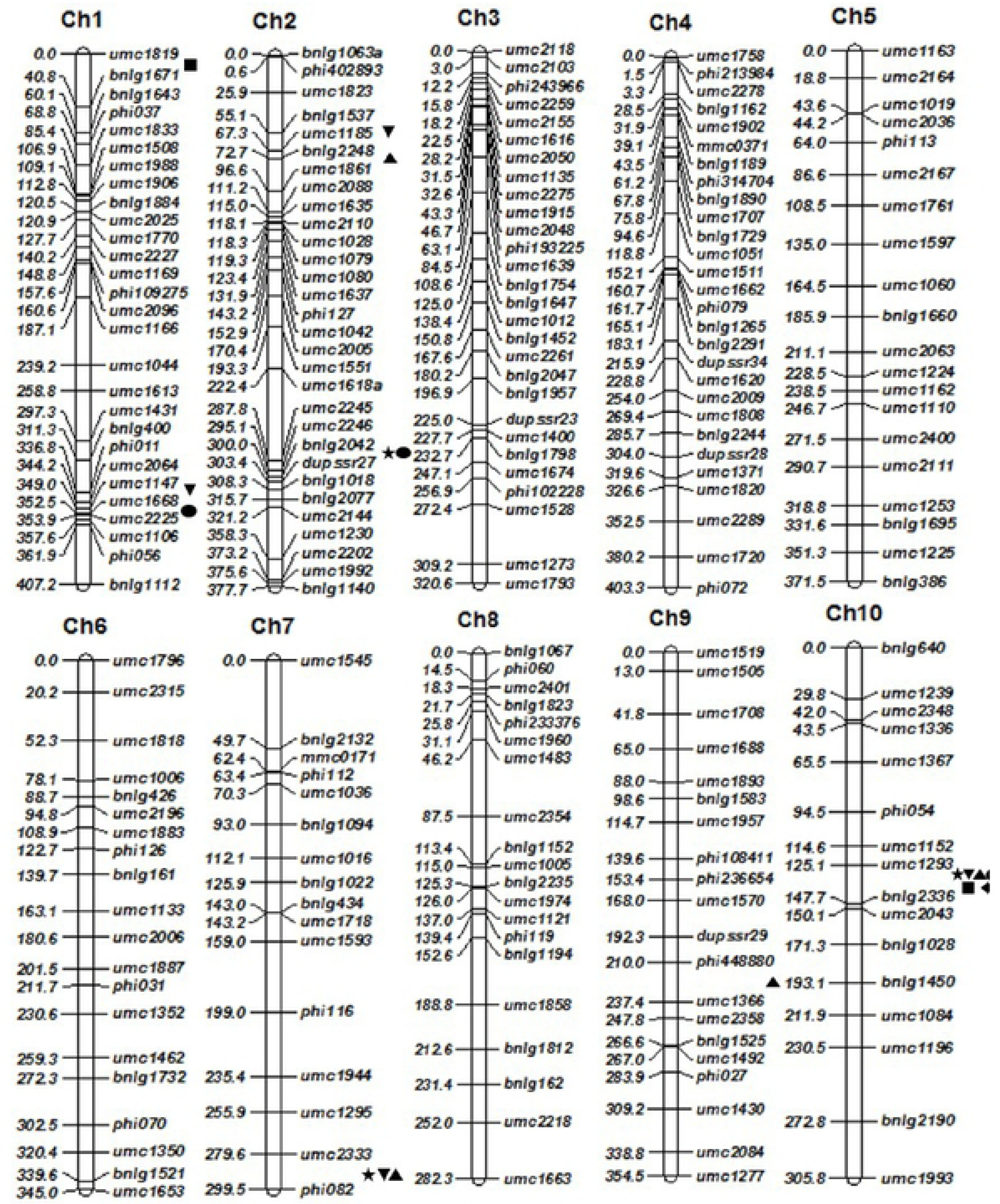
QTL for seed storability-related traits on the linkage map for RILs derived from. **★** refers to locus controlling relative germination percentage RGP; **▾** refers to locus controlling relative germination energy RGE; **▴** refers to locus controlling relative vigor index RVI; **•** refers to locus controlling relative germination energy RGE; ▪ refers to locus controlling relative seedling length RSL; ♦ refers to locus controlling relative simple vigor index RSVI

### Consistent QTL identified in the two population

A comparison of the QTL analysis results from the F2:3 and RIL populations revealed two consistent regions containing *cQTL-*7 and *cQTL-*10 on maize chromosomes 7 and 10, respectively, with positive additive effects on seed storability. *cQTL-*7, a 9.73-Mb region on chromosome 7 between markers umc1295 and umc2333, contains three QTL, *qRGP-7, qRGE-7*, and *qRGI-7*, which explain 10.52%, 12.23%, and 14.34% % of the phenotypic variation in RGP, RGE, and RGI, respectively. *cQTL-*10, a 93.13-Mb region located between the phi054 and umc2043 markers on chromosome 10, contains five QTL (*qRGP-10, qRGE-10, qRGI-10, qRVI-10*, and *qRSVI-10*), which explain 12.43%, 20.11%, 14.33%, 10.08%, and 10.85% of the phenotypic variation in RGP, RGE, RGI, RVI, and RSVI, respectively.

After adding nine and 22 SSR markers near the boundaries and internal regions of *cQTL-*7 and *cQTL-*10, respectively, QTL analyses were repeated to refine the molecular linkage maps of chromosome 7 and 10 (Figure 6). *cQTL-*7 was remapped to a 7.97-Mb region between umc1671 and phi328175 on chromosome 7 and contained four QTL, *qRGP-7, qRGE-7, qRGI-7*, and *qRSVI-7*, which explained 11.72%, 14.63%, 10.04%, and 2.27% of the phenotypic variation in RGP, RGE, RGI, and RSVI, respectively. Alleles at these QTL that increased values for traits related to seed storability came from the parental line Dong156.

**Fig. 6.**
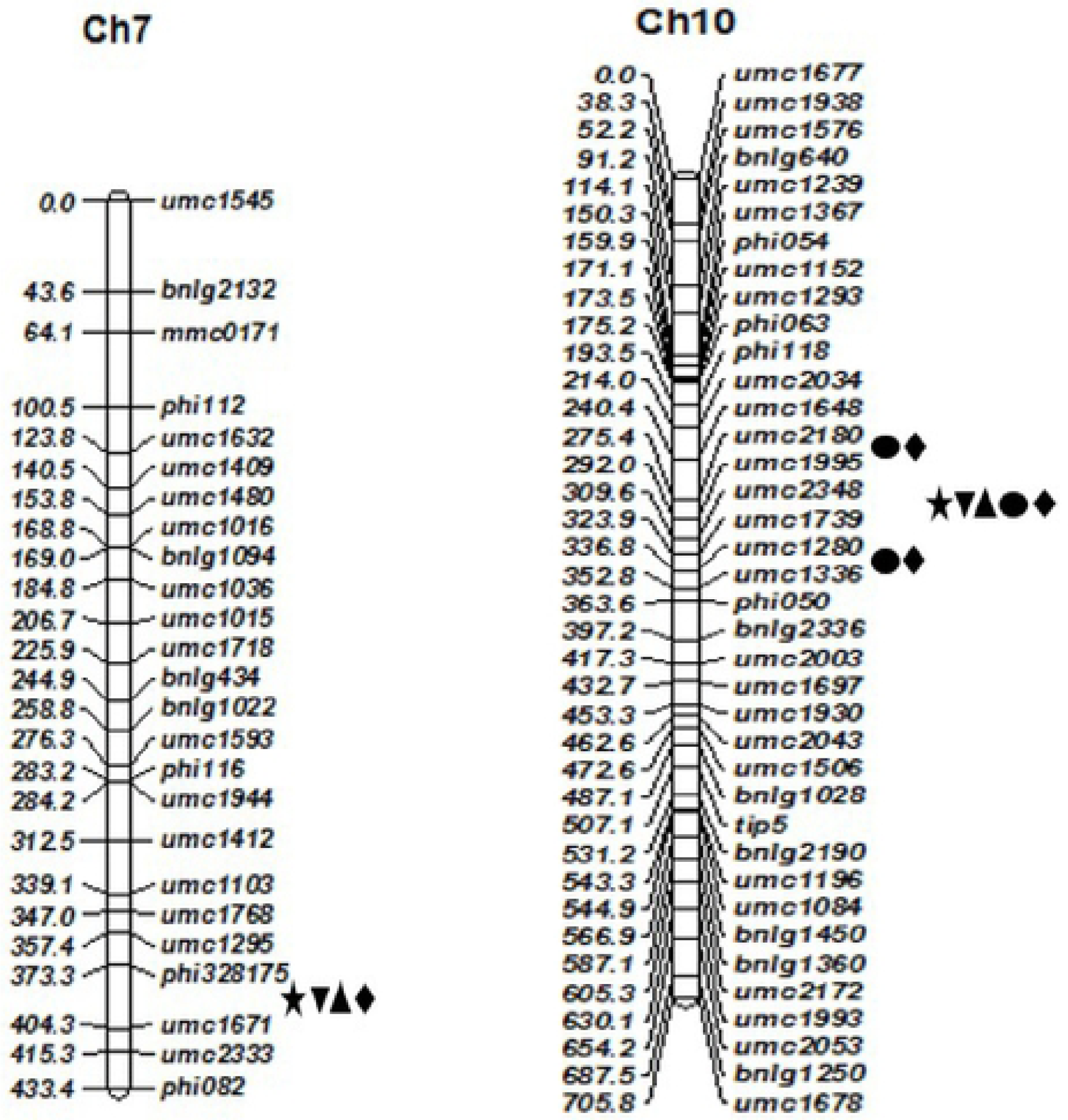
Remapping QTL for seed storability-related traits on chromosomes 7 and 10. **★** refers to locus controlling relative germination percentage (RGPO; **▾** refers to locus controlling relative germination energy (RGE); **▴** refers to locus controlling relative vigor index (RVI); **•** refers to locus controlling relative germination energy (RGE); ▪ refers to locus controlling relative seedling length (RSL); ♦ refers to locus controlling relative simple vigor index (RSVI)

*cQTL-*10 was remapped to a 39.15-Mb region on chromosome 10 between umc1648 and phi050 and contained nine QTL (*qRGP-10, qRGE-10, qRGI-10, qRVI-10-1, qRVI-10-2, qRVI-10-3, qRSVI-10-1, qRSVI-10-2* and *qRSVI-10-3*), which explained 14.19%, 17.99%, 18.26%, 22.39%, 12.80%, 19.71%, 19.59%, 19.78%, and 14.36% of the phenotypic variation in RGP, RGE, RGI, RVI, RVI, RVI, RSVI, RSVI, and RSVI, respectively (Table 3). Again, the alleles at these QTL that increased on values for traits related to seed storability all came from the parental line Dong156.

**Table 3.** Remapping of QTL for seed storability on maize chromosomes 7 and 10.

| QTL | Bin | Marker interval | Position range (cM) | LOD | Additive effect | PV (%) |
| --- | --- | --- | --- | --- | --- | --- |
| *qRGP-7* | 7.05 | *umc1671 - phi328175* | 373.3 - 404.3 | 4.01 | 0.1426 | 11.72 |
| *qRGE-7* | 7.05 | *umc1671 - phi328175* | 373.3 - 404.3 | 3.68 | 0.1418 | 14.63 |
| *qRGI-7* | 7.05 | *umc1671 - phi328175* | 373.3 - 404.3 | 3.27 | 0.1836 | 10.04 |
| *qRSVI-7* | 7.05 | *umc1671 - phi328175* | 373.3 - 404.3 | 3.43 | -0.0158 | 2.27 |
| *qRGP-10* | 10.03 | *phi054 - umc2043* | 159.9 - 462.6 | 4.03 | 0.1466 | 14.19 |
| *qRGE-10* | 10.03 | *phi054 - umc2043* | 159.9 - 462.6 | 3.89 | 0.1786 | 17.99 |
| *qRGI-10* | 10.03 | *phi054 - umc2043* | 159.9 - 462.6 | 3.85 | 0.1142 | 18.26 |
| *qRVI-10-1* | 10.03 | *phi050 - phi054* | 159.9 - 363.6 | 4.96 | 0.1418 | 22.39 |
| *qRVI-10-2* | 10.03 | *phi054 - umc2043* | 159.9 - 462.6 | 4.94 | 0.1425 | 12.80 |
| *qRVI-10-3* | 10.03 | *umc1930 - umc1648* | 240.4 - 453.3 | 4.74 | 0.1381 | 19.71 |
| *qRSVI-10-1* | 10.03 | *phi050 - phi054* | 159.9 - 363.6 | 3.92 | 0.1567 | 19.59 |
| *qRSVI-10-2* | 10.03 | *phi054 - umc2043* | 159.9 - 462.6 | 3.96 | 0.1571 | 19.78 |
| *qRSVI-10-3* | 10.03 | *umc1930 - umc1648* | 240.4 - 453.3 | 4.58 | 0.1495 | 14.36 |

### SSR makers linked to the detection and verification of major QTL

We tested nine SSR markers at the boundaries and interior regions of *cQTL-*7 and *cQTL-*10, including umc1671, phi328175, phi050, umc1648, umc1295, phi082, umc1367, phi054, and umc2043 for performing a selection efficiency test. Only the markers umc1295, phi082, umc1367, phi054, and umc2043 were polymorphic between Dong156 and Dong237. The markers umc1671, phi328175, phi050, and umc1648 showed polymorphisms between both Dong156 and Dong237 and the two resistant or susceptible pools. Individual lines from the two pools were then immediately genotyped using the above nine SSR markers. Chi-squared tests revealed a correlation between major QTL linked to seed storability. The markers umc1671, phi328175, phi050, and umc1648, with χ2 values all less than 3.84 (p = 0.05, n = 1), could thus be used for molecular marker assisted selection (MAS) for seed storability (Table 4). The selection efficiency when using these four markers was then tested in 85 RIL lines with good seed storability. On average, the correlation between genotypes and phenotypes was 83.94%, with the highest correlation between genotype and seed storability of 88.89% for the marker umc1648 (Table 5).

**Table 4.** *χ*2 tests of marker association with seed storability in pools of genotypes with good and poor storability

| Test No. | Marker | Number of plants with genotype for good storability | Number of plants with genotype for poor storability | $\chi^2$ |
| --- | --- | --- | --- | --- |
| 1 | *umc1671* | 17 | 12 | 0.862 |
| 2 | *phi328175* | 18 | 11 | 1.690 |
| 3 | *phi050* | 16 | 14 | 0.133 |
| 4 | *umc1648* | 17 | 11 | 1.286 |
| 5 | *umc1295* | 23 | 6 | 9.966 |
| 6 | *phi082* | 24 | 9 | 4.172 |
| 7 | *umc1367* | 5 | 22 | 10.704 |
| 8 | *phi054* | 19 | 8 | 4.481 |
| 9 | *umc2043* | 21 | 8 | 5.828 |

**Table 5.** Coincidence between genotype and phenotype in 85 RIL lines with good seed storability

| Marker | Number of lines |  | Degree of coincidence (%) |
| --- | --- | --- | --- |
|  | Genotype detection | Phenotype detection |  |
| *umc1671* | 64 | 85 | 83.12 |
| *phi328175* | 59 | 85 | 80.82 |
| *phi050* | 68 | 85 | 82.93 |
| *umc1648* | 72 | 85 | 88.89 |
| Average |  |  | 83.94 |

## Discussion

### Comparison of artificial seed aging methods

Sometimes during extensive natural aging of seeds, seeds can become prone to mildew and deterioration that can affect germination tests. In order to study the storability of seeds, scientists have artificially accelerated aging of plant seeds, but so far, ISTA seed evaluation procedures have only included a method for artificial aging of soybeans [25]. For other crops such as maize, there had previously been no standardized methods for artificially accelerating seed aging. Artificial aging treatments used to accelerate aging in crop seeds mainly include exposure to high temperature or high humidity [20], saturated salt [31], hot water [32], or methanol [33].

In our previous study [34], the two maize inbreds Dong156 (good seed storability) and Dong237 (poor seed storability) were used to compare the four artificial seed aging treatments mentioned above. The 58°C hot water aging treatment resulted in the greatest number of significant differences in seed storability between Dong156 and Dong237 for GP, GR, GI, VI, RGR, RGR, RGI, and RVI. The other three aging treatments did not reveal differences in seed storability between Dong156 and Dong237. Therefore, we chose a 58°C hot water treatment as the most suitable artificial aging method for maize seeds in the present study [34].

### Development of SSR markers for mining consistent QTL related to seed storability

Different QTL can be identified for a particular trait by using different genetic materials, genotypes, or phenotype detection methods. Integration and optimization of related QTL to obtain consistent QTL is very important for analyzing the genetic basis of complex traits. The physical locations of molecular markers in the genome can be determined and then used to explore other genes located within consistent QTL regions.

Studies attempting to identify consistent QTL in maize have so far focused on yield, plant type, and disease resistance, among other important traits [35-36]. For example, Semagn et al. (2013) identified 68 mQTL that could affect grain yield (GY) and anthesis silking interval (ASI) in 18 populations of maize derived from crosses between two parental lines [37]. Xu et al. (2012) surveyed the literature and collected 202 reported QTL related to flowering time and photoperiod in maize and identified 25 consensus QTL and four genomic regions containing many QTL by synthesizing the results of those reports [38]. Ali et al (2013) combined information about 389 disease-resistance QTL from 36 studies and identified a total of 10 meta-QTL for three major foliar diseases on five maize chromosomes [39]. However, few reports so far have focused on mapping of QTL and development of molecular markers related to maize seed storability [20-22].

In the present study, we compared the detection of QTL for seed storability using F2:3 and RIL populations and observed two consistent QTL that were located on separate chromosomes 7 and 10. When we remapped *cQTL-*7, which includes *qRGP-7, qRGE-7*, and *qRGI-7* between umc1671 and phi328175 (7.97Mb), we noted four previously detected QTL nearby that are related to seed vigor traits such as GE (germination energy), GP (germination percentage), GI (germination index), and VI (vigor index) and that could explain more than 10% of the phenotypic variance in these traits [40]. The QTL *qGI7a*, which influences the seed germination index [41], is close to the SSR marker phi328175 near *cQTL-*7. Further, Liu et al. (2019) [20] identified two QTL controlling GR in a 155-Mb region on chromosome 7. Han et al. (2014) [21] detected another six QTL in the same region for seed vigor-related traits in two RIL populations.

Another consistent QTL region, *cQTL-10*, which includes qRVI-10 and qRSVI-10, was remapped to a 39.15-Mb region between umc1648 to phi050 in which other QTL had previously been identified [21,40]. According the results of the present study and several previous reports, clusters of seed vigor-related QTL might reside on chromosomes 7 and 10. Compared with the results of previous reports, we refined the location of *cQTL-7* to a smaller region, which will enable further fine mapping and gene cloning.

The four SSR markers located on the boundaries of *qRGE-7, qRVI-10, umc1671, phi328175, phi050*, and *umc1648* were developed and tested in the RIL population. Seven maize inbred lines from the present study have now been designated as germplasm for improving maize seed storability by MAS and should be further verified for applied maize breeding and theoretical research.

## Acknowledgements

This research was supported by the National Key Research and Development Program China (2018YFD0100901), and by Provincial Funding for Major National Science and Technology Projects and Key Research and Development Projects (GX18B001).

## References

[1] Conelly WT, Chaiken MS. Intensive farming, agro-diversity, and food security under conditions of extreme population pressure in western Kenya. Human Ecol. 2000;28: 19–51. https://doi.org/10.1023/a:1007075621007

[2] Rajjou L, Debeaujon I. (2008). Seed longevity: Survival and maintenance of high germination ability of dry seeds. Comptes Rendus Biologies. 2008;331: 796–805. https://doi.org/10.1016/j.crvi.2008.07.021

[3] Bewley JD, Bradford K, Hilhorst H, Nonogaki H. Longevity, storage, and deterioration. In Seeds: physiology of development, germination and dormancy. 3rd ed. New York: Springer; 2013. pp. 341–376. https://doi.org/10.1007/978-1-4614-4693-4_8

[4] Bailly C. Active oxygen species and antioxidants in seed biology. Seed Sci Res. 2004;14: 93–107. https://doi.org/10.1079/ssr2004159

[5] Zacheo G, Cappello MS, Gallo A, Santino A, Cappello AR. Changes associated with post-harvest aging in almond seeds. Lebensm-Wiss U-Technol. 2000;33: 415–423. https://doi.org/10.1006/fstl.2000.0679

[6] He Y, Cheng J, Li X. Acquisition of desiccation tolerance during seed development is associated with oxidative processes in rice. Botanique. 2015;94: 91–101. https://doi.org/10.1139/cjb-2015-0154

[7] Debeaujon I, Leon-Kloosterziel KM, Koornneef M. Influence of the testa on seed dormancy, germination, and longevity in *Arabidopsis*. Plant Physiol. 2000;122(2): 403–414. https://doi.org/10.1104/pp.122.2.403

[8] Parkhey S, Naithani SC, Keshavkant S. ROS production and lipid catabolism in desiccating *Shorea robusta* seeds during aging. Plant Physiol Biochem. 2012;57: 261–267. https://doi.org/10.1016/j.plaphy.2012.06.008

[9] Xu H, Wei Y, Zhu Y, Lian L, Xie H, Cai Q, Chen Q, Lin Z, Wang Z, Xie H, Zhang J. Antisense suppression of *LOX3* gene expression in rice endosperm enhances seed longevity. Plant Biotechnol J. 2015;13: 526–539. https://doi.org/10.1111/pbi.12277

[10] Yin G, Whelan J, Wu S, Zhou J, Chen B, Chen X, Zhang J, He J, Xin X, Lu X. Comprehensive mitochondrial metabolic shift during the critical node of seed ageing in rice. PLoS One 2016;11: e0148013. https://doi.org/10.1371/journal.pone.0148013

[11] Rajjou L, Lovigny Y, Groot SP, Belghazi M, Job C, Job D. Proteome-wide characterization of seed aging in *Arabidopsis*: a comparison between artificial and natural aging protocols. Plant Physiol. 2008;148: 620–641. https://doi.org/10.1104/pp.108.123141

[12] Wu X, Liu H, Wang W, Chen S, Hu X, Li C. Proteomic analysis of seed viability in maize. Acta Physiol Plant. 2010;33: 181–191. https://doi.org/10.1007/s11738-010-0536-4

[13] Xin X, Lin XH, Zhou YC, Chen XL, Liu X, Lu XX. Proteome analysis of maize seeds: the effect of artificial ageing. Physiol Plantarum. 2011; 143:126138. https://doi.org/10.1111/j.1399-3054.2011.01497.

[14] Yan S J, Huang W J, Gao J D, Fu H, Liu J. Comparative metabolomic analysis of seed metabolites associated with seed storability in rice (*Oryza sativa* L.) during natural aging. Plant Physiol. Biochem. 2018;127: 590598. https://doi.org/10.1016/j.plaphy.2018.04.020

[15] Bettey M, Finch-Savage WE, King GJ, and Lynn JR. Quantitative genetic analysis of seed vigour and pre-emergence seedling growth traits in *Brassica oleracea*. New Phytol. 2000;148: 277–286. https://doi.org/10.1046/j.1469-8137.2000.00760.x

[16] Rehman Arif MA, Nagel M, Neumann K, Kobiljski B, Lohwasser U, Börner A. Genetic studies of seed longevity in hexaploid wheat using segregation and association mapping approaches. Euphytica. 2011;186: 1–13. https://doi.org/10.1007/s10681-011-0471-5

[17] Xie L, Tan Z, Zhou Y, Xu R, Feng L, Xing Y, Qi X. Identification and fine mapping of quantitative trait loci for seed vigor in germination and seedling establishment in rice. J Integr Plant Biol. 2014;56: 749–759. https://doi.org/10.1111/jipb.12190

[18] Zhang X, Hina A, Song SY, Kong JJ, Bhat JA, and Zhao TJ. Whole-genome mapping identified novel “QTL hotspots regions” for seed storability in soybean. (*Glycine max* L.). BMC Genomics 2019;20: 499. https://doi.org/10.1186/s12864-019-5897-5

[19] Hang NT, Lin QY, Liu LL, Liu X, Liu SJ, Wang WY, Li LF, He NQ, Liu Z, Jiang L, Wan JM. Mapping QTLs related to rice seed storability under natural and artificial aging storage conditions. Euphytica. 2015;203: 673–681. https://doi.org/10.1007/s10681-014-1304-0

[20] Liu YN, Zhang HW, Li XH, Wang F, Lyle D, Sun LJ, Wang GY, Wang JH, Li L, Gu RL. Quantitative trait locus mapping for seed artificial aging traits using an F2:3 population and a recombinant inbred line population crossed from two highly related maize inbreds. Plant Breeding. 2019;138: 29–37. https://doi.org/10.1111/pbr.12663

[21] Han Z, Bin W, Zhang J, Guo S, Zhang H, Xu L, and Chen Y. Mapping of QTLs associated with seed vigor to artificial aging using two RIL populations in maize (*Zea mays* L.). Agricultural Sci. 2018;9: 397–415. https://doi.org/10.4236/as.2018.94028

[22] Han Z, Ku L, Zhang Z, Zhang J, Guo S, Liu H, Zhao R, Ren Z, Zhang L, Su H, Dong L, Chen Y. QTLs for seed vigor related traits identified in maize seeds germinated under artificial aging conditions. PLoS One. 2014;9: e92535. https://doi.org/10.1371/journal.pone.0092535

[23] Murray MG, Thompson WF. Rapid isolation of high molecular weight plant DNA. Nucleic Acids Res. 1980;8: 4321–4326. https://doi.org/10.1093/nar/8.19.4321

[24] Di H, Liu XJ, Wang QK, Weng JF, Zhang L, Li XH, Wang ZH. Development of SNP-based dCAPS markers linked to major head smut resistance quantitative trait locus qHS2.09 in maize. Euphytica. 2015;202: 69–79. https://doi.org/10.1007/s10681-014-1219-9

[25] Chapter 5: the germination test. In: International Rules for Seed Testing. Bassersdorf, Switzerland: International Seed Testing Association (ISTA); 2015. pp. i-5–56. https://doi.org/10.15258/istarules.2019.05

[26] Kishor DS, Lee C, Lee D, Venkatesh J, Seo J, Chin JH, Jin Z, Hong SK, Ham JK, Koh HJ. D Novel allelic variant of *Lpa1* gene associated with a significant reduction in seed phytic acid content in rice (*Oryza sativa* L.). PLoS One. 2019;14(3): e0209636. https://doi.org/10.1371/journal.pone.0209636

[27] Xi XJ, Zhu YG, Tong YP, Yang XL, Tang NN, Ma SM, Li S, Cheng Z. Assessment of the genetic diversity of different Job’s Tears (*Coix lacryma-jobi* L.) Accessions and the active composition and anticancer effect of its seed oil. PLoS One. 2016;11(4): e0153269. https://doi:10.1371/journal.pone.0153269

[28] Zhang Y, Huo M, Zhou J, Xie S. PKSolver: an add-in program for pharmacokinetic and pharmacodynamic data analysis in Microsoft Excel. Comput Methods Prog Biomed. 2010;99: 306–14. https://doi.org/10.1016/j.cmpb.2010.01.007

[29] Lander ES, Green P, Abrahamson J, & Barlow, A. (1987). MAPMAKER: An interactive computer package for constructing primary genetic linkage maps of experimental and natural populations. Genomics, 1, 174–181. https://doi.org/10.1016/0888-7543(87)90010-3

[30] Li XH, Wang ZH, Gao SR, Shi H L, Zhang SH, George MLC, Li MS, and Xie CX. Analysis of QTL for resistance to head smut (*Sporisorium reiliana*) in maize. Field Crops Res. 2008;106: 148–155. https://doi.org/10.1016/j.fcr.2007.11.008

[31] Bennett MA, Grassbaugh EM, Evans AF, Kleinhenz MD. Saturated salt accelerated aging (SSAA) and other vigor tests for vegetable seeds. Seed Technol. 2004; 26: 67–74. https://doi.org/10.2307/23433494

[32] Kodde J, Buckley WT, de Groot CC, Retiere M, Zamora AMV, Groot SPC. A fast ethanol assay to detect seed deterioration. Seed Sci Res. 2012;22: 55–62. https://doi.org/10.1017/S0960258511000274

[33] Addai IK, Safo-Kantanka O. Evaluation of screening methods for improved storability of soybean seed. Intl J Bot. 2006;2: 152–155. https://doi.org/10.3923/ijb.2006.152.155

[34] Di H, Lv TT, Liu LL, Xia HM, and Wang ZH. Comparative analysis on phenotypic measurement of maize seeds storability by artificial aging methods. J Maize Sci. 2015;23: 71–75. (in Chinese with English abstract) https://doi.org/10.13597/j.cnki.maize.science.20150312

[35] Wang YJ, Xu J, Deng DX, Ding HD, Bian YL, Yin ZT, Wu YR, Zhou B, Zhao Y. A comprehensive meta-analysis of plant morphology, yield, stay-green, and virus disease resistance QTL in maize (*Zea mays* L.). Planta. (2016);243: 459–471. https://doi.org/10.1007/s00425-015-2419-9

[36] Wang YJ, Wang YL, Wang X, Deng DX. Integrated meta-QTL and genome-wide association study analyses reveal candidate genes for maize yield. J Plant Growth Regul. 2019; pp. 1–10. https://doi.org/10.1007/s00344-019-09977-y

[37] Semagn K, Beyene Y, Warburton ML, Tarekegne A, Mugo S, Meisel B, Sehabiague P, and Prasanna BM. Meta-analyses of QTL for grain yield and anthesis silking interval in 18 maize populations evaluated under water-stressed and well-watered environments. BMC Genomics. 2013;14: 313. https://doi.org/10.1186/1471-2164-14-313

[38] Xu J, Liu YX, Liu J, Cao MJ, Wang J, Lan H, Xu YB, Lu YL, Pan GT, Rong TZ. The genetic architecture of flowering time and photoperiod sensitivity in maize as revealed by QTL review and meta analysis. J Integr Plant Biol. 2012;54: 358–373. https://doi.org/10.1111/j.1744-7909.2012.01128.x

[39] Ali F, Pan Q, Chen G, Zahid KR, Yan J. Evidence of multiple disease resistance (MDR) and implication of meta-analysis in marker assisted selection. Plos One. 2013;8: e68150. https://doi.org/10.1371/journal.pone.0068150

[40] Liu JB, Fu ZY, Xie HL, Hu YM, Liu ZH, Duan LJ, Xu SZ, Tang JH. Identification of QTLs for maize seed vigor at three stages of seed maturity using a RIL population. Euphytica. 2011;178: 127–135. https://doi.org/10.1007/s10681-010-0282-0

[41] Wang B, Zhang ZH, Fu ZY, Liu ZH, Hu YM, Tang JH. Comparative QTL analysis of maize seed artificial aging between an immortalized F2 population and its corresponding RILs. Crop J. 2016;4: 30–39. https://doi.org/10.1016/j.cj.2015.07.004

